# 14K Prolactin Derived 14-Mer Antiangiogenic Peptide Targets Bradykinin-/Nitric Oxide-cGMP-Dependent Angiogenesis

**DOI:** 10.1101/2023.08.21.554080

**Authors:** Jaeok Lee, Pavitra Kumar Jadaun, Suganya Natarajan, So Hyeon Park, Syamantak Majumder, Lakshmikirupa Sundaresan, Kambadur Muralidhar, Jong-Soon Choi, Hwa Jeong Lee, Suvro Chatterjee

## Abstract

VEGF-targeted antiangiogenic therapy for cancers has been principally used but also faced a limitation due to resistance and adverse effects in clinical application. This observation further endorses the need for novel anti-angiogenesis molecules and/or understanding of the mechanisms of tumor angiogenesis before clinical trial. In the present study, we investigated the antiangiogenic properties of a novel 14-mer antiangiogenic peptide (14-MAP) derived from N-terminal 14kDa buffalo prolactin, followed by an exploration of its mode of action. 14-MAP at the picomolar concentration inhibited VEGF- and bradykinin (an autacoid peptide expressed in vascular tissues in pathophysiology)-stimulated endothelial nitric oxide (eNO) production, cell migration and proliferation in endothelial cells and vessel development in chick embryo. The crucial inhibitory effects of the peptide, however, were presented on the bradykinin-dependent angiogenic properties. Moreover, the interference of 14-MAP with the eNO synthase (eNOS)-cyclic GMP pathway was identified. A combination of low dose of Avastin, a widely used drug targeting VEGF-dependent angiogenesis, and 14-MAP significantly reduced tumor size in a mouse model of human colon cancer. These results suggest that 14-MAP, a bradykinin- and eNOS-dependent antiangiogenic peptide, can be useful for overcoming the limitation of VEGF-targeted antiangiogenic therapy in cancer patients.

## INTRODUCTION

Angiogenesis is the process of development of new blood vessels from preexisting vasculature. The process is fundamental and critical to physiological and pathological developments (1-3). In 1865, Rudolf Virchow observed a huge vascular network in growing tumors and the observation has been followed for the next 150 years until Judah Folkman introduced the concept of anti-angiogenesis therapy for perfused tumors by targeting and attenuating a critical growth factor, the vascular endothelial growth factor (VEGF) (4-5). For last decades, the anti-angiogenesis targeting VEGF in cancer and metastasis has been studied for leading to the approval in clinic (6). Although the VEGF-targeted antiangiogenic approaches have been used for cancer therapy, the application faces a limitation; presently Avastin, the first antiangiogenic drug, is approved to treat metastatic colorectal cancer for first-or second-line treatment in combination with fluorouracil-based chemotherapy, but not approved for using after surgery (5,7). Therefore, in-depth mechanistic insight into anti-angiogenesis therapy that considers other hallmarks of the tumor is required (5). The search for better angiogenic candidates, which target other critical growth factors and natural peptides, is back on track. Our previous study introduced 14kDa (K) buffalo prolactin (buPRL) which antagonized VEGF- and bradykinin (BK)-dependent endothelial nitric oxide (eNO) production, essential for cell migration and tube formation (8). We synthesized a 14-mer antiangiogenic peptide (14-MAP) sequenced in the N-terminal 14K buPRL (8).

BK, the classical 9-mer peptide produced by the kallikrein-kinin system, is implicated in pathophysiological processes such as inflammation, cell proliferation, cell migration, coagulation, fibrogenesis, angiogenesis and cancer (9-12). BK is an agonist for endothelial NO synthesis by promoting sub-cellular trafficking of eNOS in endothelial cells (13). In addition, the BK receptors, B1R and B2R are known to play an important role in cancer growth and metastasis. Elevated expression of B1R and B2R have been observed in many cancers including colon cancer (12-18). Therefore, targeting BK-receptor pathway will be useful for cancer therapy.

The endothelial NO synthase (eNOS) pathway is essential process for angiogenesis and vascular permeability and relaxation stimulated by vasoactive factors, including VEGF, BK and eNOS-produced NO production is positively correlated with endothelial cell migration (19,20). The vascular eNOS/NO/soluble guanylatecyclase (sGC)/cyclic GMP (cGMP)/cGMP-dependent protein kinase (PKG) pathway is activated by PI3K/Akt signaling in the endothelium (21). Gonzalez et al. reported that the activation of eNOS mediated by VEGF, BK and acetylcholine (Ach) was inhibited by N-terminal 16K human PRL (hPRL) in bovine and human umbilical vein endothelial cells (BUVEC and HUVEC) (22). We observed, however, that the inhibition of eNO production by 14K buPRL in the immortalized EAhy926 cells was related with VEGF and BK only, but not with Ach (8). BK, the agonist of eNOS plays a critical role in the sub-cellular localization of eNOS (13) along with its involvement in the known BK-PI3K-Calcium calmodulin-eNOS signaling pathway (23).

In the present study, we demonstrated strong inhibitory effects of 14-MAP in picomolar concentrations on BK-induced endothelial functions like NO production, cell migration and chick embryonic vessel formation. Moreover, the mechanistic insight of antiangiogenic properties of the peptide was deciphered and the enhancement effect on the reduction of human colon tumor size of 14-MAP was confirmed when co-treated with a low dose of Avastin.

## MATERIALS AND METHODS

### Materials

The immortalized human endothelial cell line, EAhy926 and the human colon adenocarcinoma cell line, HT29 were gifted from Dr. C.J.S. Edgel (University of North Carolina, USA) and Dr. U. Schumacher (University of Medical Center Hamburg-Eppendorf, Germany), respectively. Brown leghorn (*Gallus gallus*) fertilized eggs and Balb/c nude mice (*nu/nu*) were purchased from a poultry farm (Potheri, Chennai, India) and Orient Bio (Seongnam, Korea), respectively.

14-MAP (AQGKGFITMALNSC (pI 8.27/molecular weight (M.W.) approx.1.4kDa) and 14-MAP scramble (Scr) (GSQCAAGTMNLKIF (http://users.umassmed.edu/ian.york/Scramble.shtml accessed on 7^th^ Jan. 2015)) were gifted from Dr. J.K. Bang (Korea Basic Science Institute (KBSI), Korea). BK, VEGF, des-Arg9-BK (B1R agonist), des-Arg9-Leu8]-BK (B1R antagonist) and 3-(5′-Hydroxymethyl-2′-furyl)-1-benzyl indazole were purchased from Sigma-Aldrich (St. Louis, MO, USA). 8-Bromoguanosine 3′, 5′-cyclic monophosphate and DAF-FM were purchased from Calbiochem (Darmstadt, Germany) and Molecular Probes^®^ (Thermo Fisher Scientific, Rockford, IL, USA), respectively. Avastin was obtained from Selleckchem (Houston, TX, USA). SpermineNONOate was purchased from Cayman Chemicals (Ann Arbor, MI, USA). All other reagents were of molecular or cell culture grade.

### Cell Culture

EAhy926 cells (24) were cultured in DMEM supplemented with 10% fetal bovine serum and penicillin (100 U/mL)/streptomycin (50 pg/mL) at 37ºC and 5% CO_2_. For the xenograft model, HT29 cells were maintained in RPMI with the same conditions as mentioned above.

### Scrape Wound Healing Assay

The endothelial cells were cultured as a monolayer in a 96-well plate up to 90% confluence. One straight-line scratch was made across each well using a sterile p10 pipet tip as described (25). Debris from the scratch area was washed with media. Cells were treated with the media with 10% FBS containing the respective treatment and incubated. Bright-field images were taken under 4X magnification with a camera adapted to the Olympus XL70 bright field microscope at every 2 h interval. HT29 cells were cultured as a monolayer in a 24-well plate up to 85-90% confluence. Cells were treated with the media without serum, containing the respective treatment. Images were taken under 5X magnification with a camera adapted to Zeiss AXIO observer 7 inverted microscope (Axio observer 7, Car Zeiss, Jena, Germany) equipped at Drug Development Research Core Center in Ewha Womans University, at every 24 h interval. Finally, the percentage of wound healing was quantified from the images using ImageJ software. The data was collected and plotted from a minimum of three independent experiments. The assay was normalized for cell number, viability and proliferation index.

### NO Production Assay

NO was measured using DAF-FM fluorescence probe. EAhy926 cells were seeded in a 96-well plate and incubated overnight to obtain 90-95% confluence. The media was replaced with appropriate treatment media and incubated for 2 h in a CO_2_ incubator. After incubation the media was removed, cells were washed with PBS and 100 μL of DAF-FM (3 μM) in PBS was added to each well. The plate was incubated for 20 min in dark. 100 μL of supernatant from cells was collected and transferred to a fresh 96-well plate. Then absorbance was read at 495/515 nm using a fluorescence spectrophotometer (Cary Eclipse, Agilent technology, Bangalore, India). The data was collected and plotted from a minimum of three independent experiments.

### Cell Proliferation Assay

The endothelial cells were seeded in a 96-well plate around 15-20% confluency and incubated for 24 h. After starvation for 1-18 h, cells were treated with each factor with 0.2% serum medium, and then an MTT assay (26) was performed on the cells 48 h later. The absorbance was read at 540 nm using a microplate reader (Bio-Rad, 680). Cell viability was expressed as percentage of cell proliferation. HT29 cells were seeded in a 96-well plate around 15-20% confluency and incubated for 24 h. Cells were treated with each factor without serum medium, and then sulforhodamine B staining assay (27) was performed on the cells 24 and 48 h later. The data was collected and plotted from a minimum of three independent experiments.

### Chick Chorioallantoic Membrane (CAM) Assay

CAM assay was performed with fertilized eggs at Hamburger-Hamilton’s stage 16 of chick development (25,28). The eggs were incubated at 37°C in a humidified incubator (Southern Incubators, Chennai, India) for 3 days and on the fourth day opened north polar under sterile conditions. Whatman filter paper disks contacting target factors including control were placed over the vasculature of area vasculosa, and then the eggs were incubated for 8 h. The vasculature was imaged every two hours from the time of drug treatment. All images were quantified using the Angio-Quant software (29) and total vessel length, number of junctions and vessels size were calculated. The percentage change in these parameters was calculated and plotted against the treatment. The data was collected and plotted from a minimum of three independent experiments.

### Xenograft Nude Mice Experiment

HT29 (3.5-5.0 × 10^6^ cells/100 μL PBS and Matrigel^®^ mixture) was subcutaneously inoculated on the flank of Balb/c nude mice (5-week-old (18.2-22.5 g), male). When tumor volume was approximately 100 mm^3^, the treatments of 14-MAP, Avastin and the combination of both versus control (PBS) were performed every 3 days for 3 weeks by an intravenous injection to the tail vein (*n* = 5/group). The body weight was monitored, and tumor size was measured with a digital caliper twice a week. The tumor volume was calculated using the formula (length x width^2^)/2. All animal care and experimentation followed the procedure approved by the Ewha Womans University Institutional Animal Care and Use Committee (No. 2016-062), Korea.

### Statistical Analysis

Statistical analysis was conducted using Tukey’s or Dunnett’s test in conjunction with one-way ANOVA test or Dunnett’s T3 test in conjunction with Welch one-way ANOVA. For in vivo experiments, Student’s *t*-test and nonparametric Kruskal-Wallis test were performed. GraphPad Prism was used for the analysis (Version 8.0.0 (224), La Jolla, CA, USA). All mean data were represented with standard error (SEM). The *P* value < 0.05 was set as a statistical significance.

## RESULTS

### 14-MAP Inhibits VEGF- and BK-Induced EC Functions

The antiangiogenic effect of the 14-MAP peptide was confirmed by wound healing analysis (Figure 1A and B). Excessively low concentration of the 14-MAP peptide sufficiently inhibited EC migration. The significant inhibition was progressive in a time-dependent manner, compared to the control (*P* < 0.05-0.01). Scr peptide did not show any difference from the control. The inhibitory effect was validated in chick embryo vessel development (Figure S1). The working concentration of 14-MAP was determined in the range of pico-molarity and had no cytotoxicity (Figure S2).

**Figure 1.**
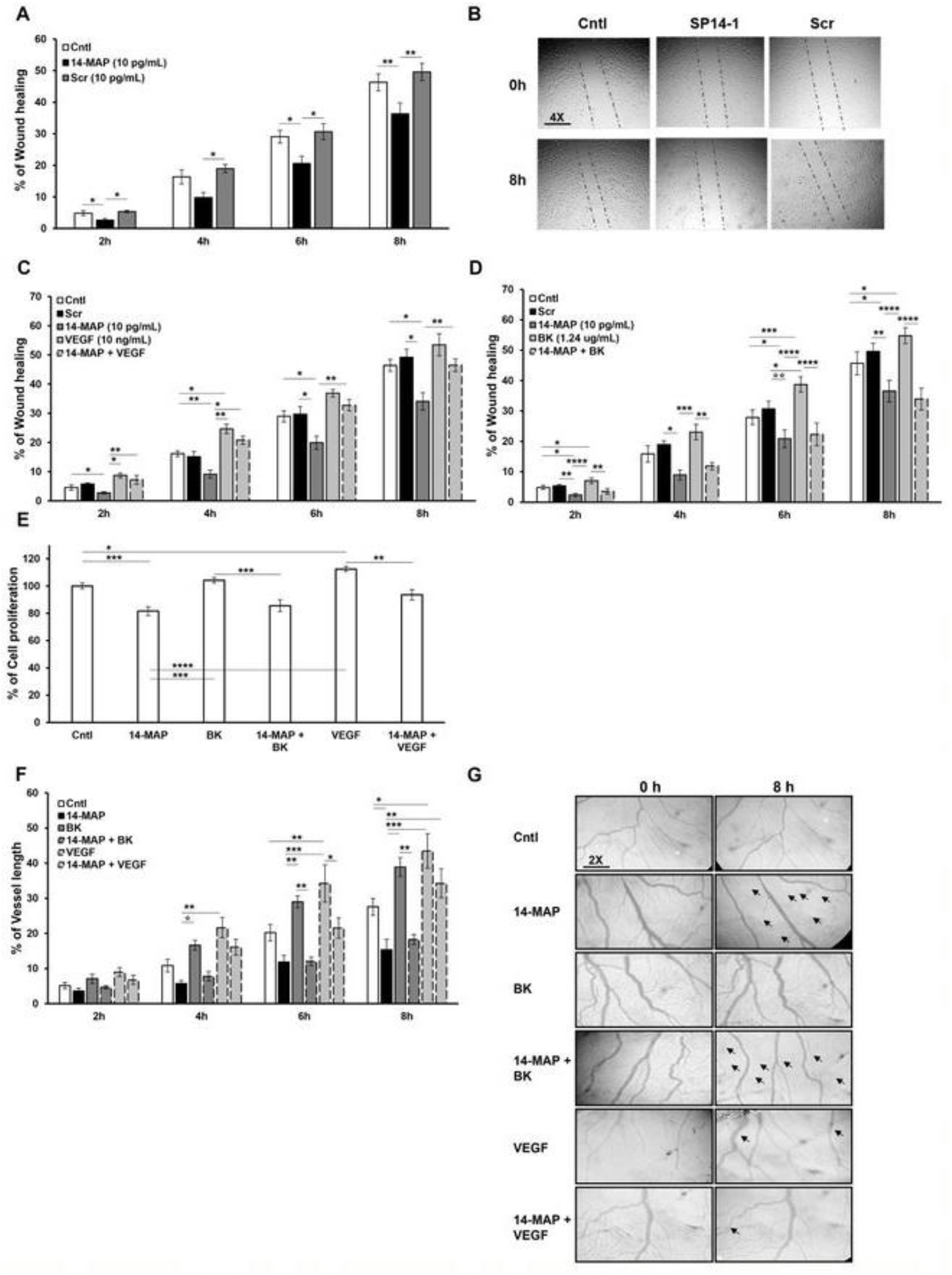
Inhibition of VEGF- and BK-induced endothelial cell functions by 14-MAP. **A** and **B**, Cell migration analysis by wound healing assay. Cell migration analysis by Wound healing assay of 14-MAP on VEGF pathway (**C**) and on BK pathway (**D**) **E**, Cell proliferation analysis, and **F** and **G**, Angiogenic vessel growth analysis by CAM assay of 14-MAP on VEGF- and BK-pathways, respectively. 14-MAP and Scr, 10 pg/mL (approx. 7 pM); VEGF, 10 ng/mL (approx. 0.3 nM); BK, 1.24 μg/mL (approx. 1.24 μM). Cntl, control; Scr, scramble (GSQCAAGTMNLKIF). *, *P* < 0.05; **, *P* < 0.01; ***, *P* < 0.001; ****, *P* < 0.0001. Arrows represent indicate the deteriorating (14-MAP, 14- MAP+BK and 14-MAP+VEGF) and ameliorating (VEGF) vascular network (**G**).

To understand the 14-MAP-mediated antiangiogenic pathway, we investigated the effects of the peptide on VEGF- and BK-induced EC functions, confirmed as related pathways to the inhibition of 14K buPRL in our previous study (8). 14-MAP inhibited the cell migration stimulated by exogenous VEGF and BK as well as endogenous EC migration significantly (*P* < 0.05) (Figure 1C and D). The inhibition value of 14-MAP on the exogenous BK-increased EC migration developed in a time-dependent manner (*P* < 0.01-0.0001) and was similar to that of the peptide on the endogenous EC migration (*P* < 0.05) (Figure 1D), whereas the inhibition level of 14-MAP on the exogenous VEGF-increased EC migration was just similar with the control level (Figure 1C). Scr peptide did not significantly affect BK-induced EC migration (Figure S3A).

Moreover, the inhibitory effect of 14-MAP on VEGF- and BK-increased EC proliferation (for 48 h) was investigated (Figure 1E). 14-MAP significantly inhibited EC proliferation up to approximately 20% compared to the control (*P*< 0.001). Exogenous VEGF- and BK-induced EC proliferation appeared to increase 5 and 10%, respectively, compared to the control. The exogenously stimulated EC proliferations were significantly inhibited by 14-MAP (*P* < 0.001 and *P* < 0.01, respectively) as much as endogenous inhibition (approximately 20%) by the peptide. Scr did not significantly affect EC proliferation or VEGF- and BK-induced proliferation, either (Figure S3B).

14-MAP inhibition of VEGF- and BK-induced EC functions estimated by an *in vitro* system was confirmed by an *ex vivo* system. The embryo vessel development enhanced by VEGF or BK was inhibited by the antiangiogenic peptide, 14-MAP (Figure 1F and G, Figure S3C and D). BK-enhanced vessel length development was decreased by 14-MAP (*P* < 0.01) as much as 14-MAP-decreased vessel length development (*P* < 0.05), while VEGF-enhanced vessel length development (*P* < 0.05-0.01) was decreased by 14-MAP as much as the control level (Figure 1F and G). These patterns were also monitored in the development of embryo vessel size and junctions (Figure S3C and D). These results of *in vitro* and *ex vivo* tests indicate that the antiangiogenic peptide 14-MAP is more correlated to the BK pathway than the VEGF pathway on EC functions.

### 14-MAP Interferes with eNOS Pathway on EC Migration

In order to reveal the mechanism of 14-MAP interference on angiogenesis, we investigated the effect of 14-MAP on the eNOS pathway. 14-MAP-mediated NO inhibition was confirmed on endogenous NO production in EC (approximately 0.5 RFU value decrease (*P* < 0.01) *vs* control) (Figure 2A). Moreover, the peptide decreased exogenous BK- and VEGF-enhanced NO production (0.8 decrease (*P* < 0.001) and 0.8 decrease (*P* < 0.001), respectively) (Figure 2A). Scr peptide did not induce any significant changes in the NO production increased by BK and VEGF as well as in the NO level of control.

**Figure 2.**
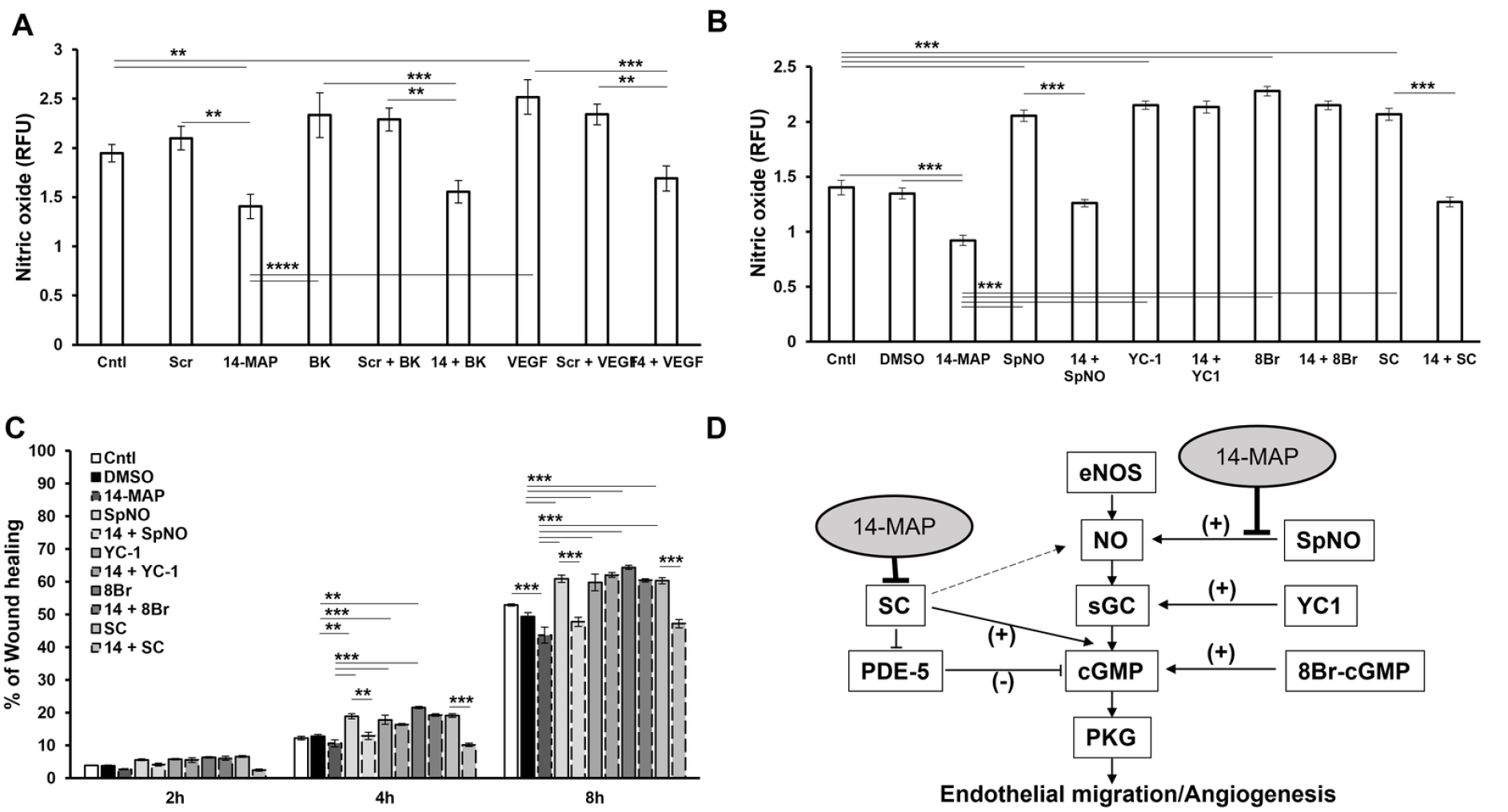
Inhibitory effect of 14-MAP in eNOS pathway. **A**, NO production analysis of 14-MAP on VEGF pathway in EC. **B**, NO production analysis of 14-MAP on eNOS/NO/sGC/cGMP pathway in EC. **C**, Cell migration analysis of 14-MAP on eNOS/NO/sGC/cGMP pathway in EC. **D**, Diagram of eNOS/NO/sGC/cGMP pathway correlated with 14-MAP. Cntl, control; Scr, scramble (GSQCAAGTMNKIF). **, *P* < 0.01; ***, *P* < 0.001; ****, *P* < 0.0001.

To comprehend what step in the pathway is interfered by 14-MAP on EC migration, we also studied NO production in EC (Figure 2B) and EC migration (Figure 2C) after co-treatment of 14-MAP with each inducer, SpNO (NO donor) (30), YC1 (soluble guanylate cyclase inducer) (31), 8Br-cGMP (cGMP analog) (32) and SC (cGMP stabilizer) (33), on the eNOS/NO/sGC/cGMP pathway (Figure 2D). DMSO used for dissolving the NO inducers did not affect NO production and EC migration (Figure 2B and C). 14-MAP significantly inhibited NO production increased by SpNO (0.7, *P* < 0.001) and SC (0.8, *P* < 0.001), but not by YC1 and 8Br-cGMP (Figure 2B). The inhibited level by 14-MAP on the SpNO- and SC-increased NO was as much as the NO production of the control. This pattern was confirmed on EC migration (Figure 2C). 14-MAP inhibited EC migration induced by SpNO (13.1% at 8 h, *P* < 0.001) and SC (13.1% at 8 h, *P* < 0.001) time dependently, but no differences were monitored on YC- and 8 Br-cGMP-induced EC migration when co-treated with 14-MAP (Figure 2C). Therefore, it was observed that SpNO- and SC-induced NO production and EC migration were correlated with 14-MAP inhibition (Figure 2D).

### 14-MAP Interferes with BK-BK Receptor Pathway on EC Functions

Meanwhile, we considered to the correlation between 14-MAP and BK pathway (Figure 1D, F and G). To understand which BK receptor (B1R or B2R) (Figure S4) was related to 14-MAP, we investigated the effect of 14-MAP on BK-B1R and -B2R pathways by EC cell migration, proliferation, and NO production (Figure 3). When 14-MAP was treated with an agonist of B1R and B2R in EC, the peptide significantly inhibited Des-Arg9-BK (R9, B1R agonist)- and BK (B2R agonist)-increased EC migrations (12% (*P* < 0.05) and 21% (*P* < 0.0001) at 8 h, respectively) as well as an endogenous EC migration (12% (*P* < 0.05) at 8 h) (Figure 3A). B1R antagonist [Leu8]Des-Arg9-BK (L8) also inhibited R9- and BK-increased EC migrations (17% (*P* < 0.001) and 13% (*P* < 0.01) at 8 h, respectively) as well as endogenous EC migration (8% at 8 h) (Figure 3A). In addition, the combination treatment of 14-MAP with L8 induced a similar level of inhibition as by 14-MAP alone, compared to control (14% (*P* < 0.001) at 8 h). The level of 14-MAP inhibition of the B2R agonist (BK)/B2R pathway was the highest, and the level of 14-MAP inhibition of the B1R agonist (R9)/B1R pathway was similar to the level of B1R antagonist (L8) inhibition of the B2R agonist/B2R pathway. Therefore, the inhibition effect of 14-MAP showed more interference to BK/B2R pathway-induced EC migration.

**Figure 3.**
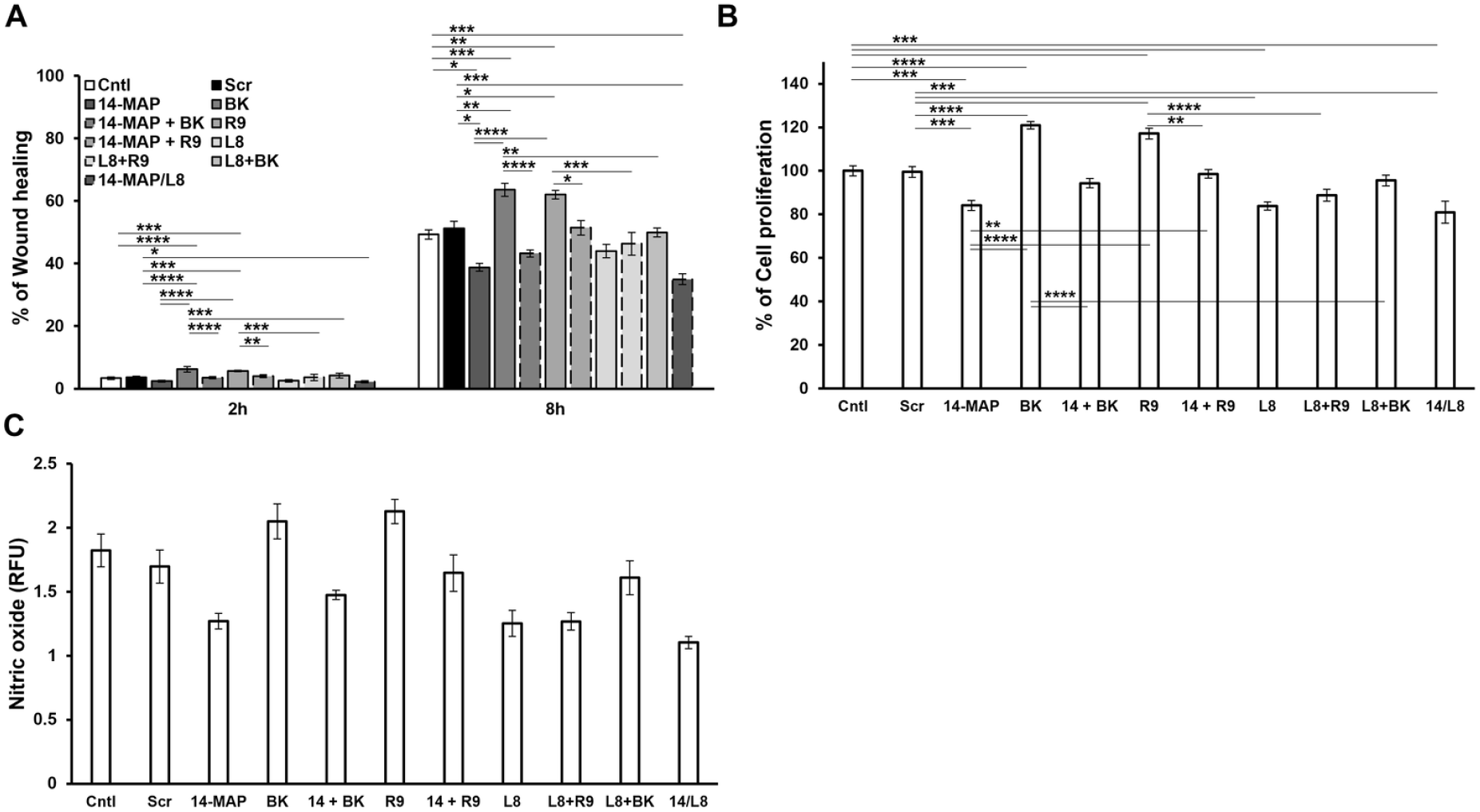
Interference of BK-BKRs pathway by 14-MAP. **A**, EC cell migration. **B**, EC cell proliferation. **C**, NO production. Cntl, control; Scr, scramble; BK, B2R agonist; 14, 14-MAP; R9, Des-Arg9-BK/B1R agonist; L8, [Leu8]Des-Arg9-BK/B1R antagonist. *, *P* < 0.05; **, *P* < 0.01; ***, *P* < 0.001; ****, *P* < 0.0001.

This pattern was also confirmed on EC proliferation (Figure 3B) and eNO production (Figure 3C). 14-MAP significantly inhibited exogenous BK- and R9-induced EC proliferation increase (24% (*P* < 0.0001) and 18% (*P* < 0.01), respectively) as well as endogenous EC proliferation (16%, *P* < 0.001), and L8 inhibited exogenous BK- and R9-increased EC proliferation (25% (*P* < 0.0001) and 28% (*P* < 0.0001), respectively) as well as endogenous EC proliferation by L8 (16%, *P* < 0.001) (Figure 3B). The combination treatment of 14-MAP with L8 was similar to the inhibition level of 14-MAP alone or L8 alone, compared to control (14%, *P* < 0.001). The inhibition of BK- and R9-increased eNO production as well as endogenous eNO production by 14-MAP (0.6%, 0.5% and 0.5%, respectively), and BK- and R9-increased eNO productions as well as endogenous eNO production by L8 (0.5%, 0.8% and 0.6%, respectively) was confirmed (Figure 3C). The combination treatment of 14-MAP with L8 was slightly higher than the inhibition level of 14-MAP alone or L8 alone compared to control (0.7%) (Figure 3C). Finally, these results led to the idea that 14-MAP-mediated inhibition was related to the BK-receptor pathway, and more interfered to BK/B2R pathway than R9/B1R pathway on EC functions.

### The Antiangiogenic Peptide 14-MAP Inhibits Human Colon Cancer Growth *In Vivo*

Furthermore, we investigated the effect of the antiangiogenic peptide in cancer (Figure 4). The low dose of 14-MAP (0.5 μg/kg), not toxic to mice (Table S1 and Figure S5) was given to human colon cancer cells (HT29)-xenograft nude mice by intravenous injection (Figure 4A and B). The effect of the peptide on reduced tumor growth compared to control was monitored and found to be less than the effect of Avastin, a well-known anticancer agent targeting VEGF (Figure 4A). The low dose of Avastin (0.5 mg/kg) was used in the present study (Figure S5A). The reduction of tumor growth was more significantly enhanced by co-treatment of 14-MAP with Avastin (*P* < 0.05-0.01) (Figure 4A). The statistical difference was between the groups of the control and the combination treatment only, among each group (Figure 4A). No treatment induced any change in body weight in the mice (Figure 4B). However, the pattern of the change in the treatment period showed that the combination of 14-MAP with Avastin was similar to the control, and 14-MAP only and Avastin only were slightly less than control at the latter half period (Figure 4B).

**Figure 4.**
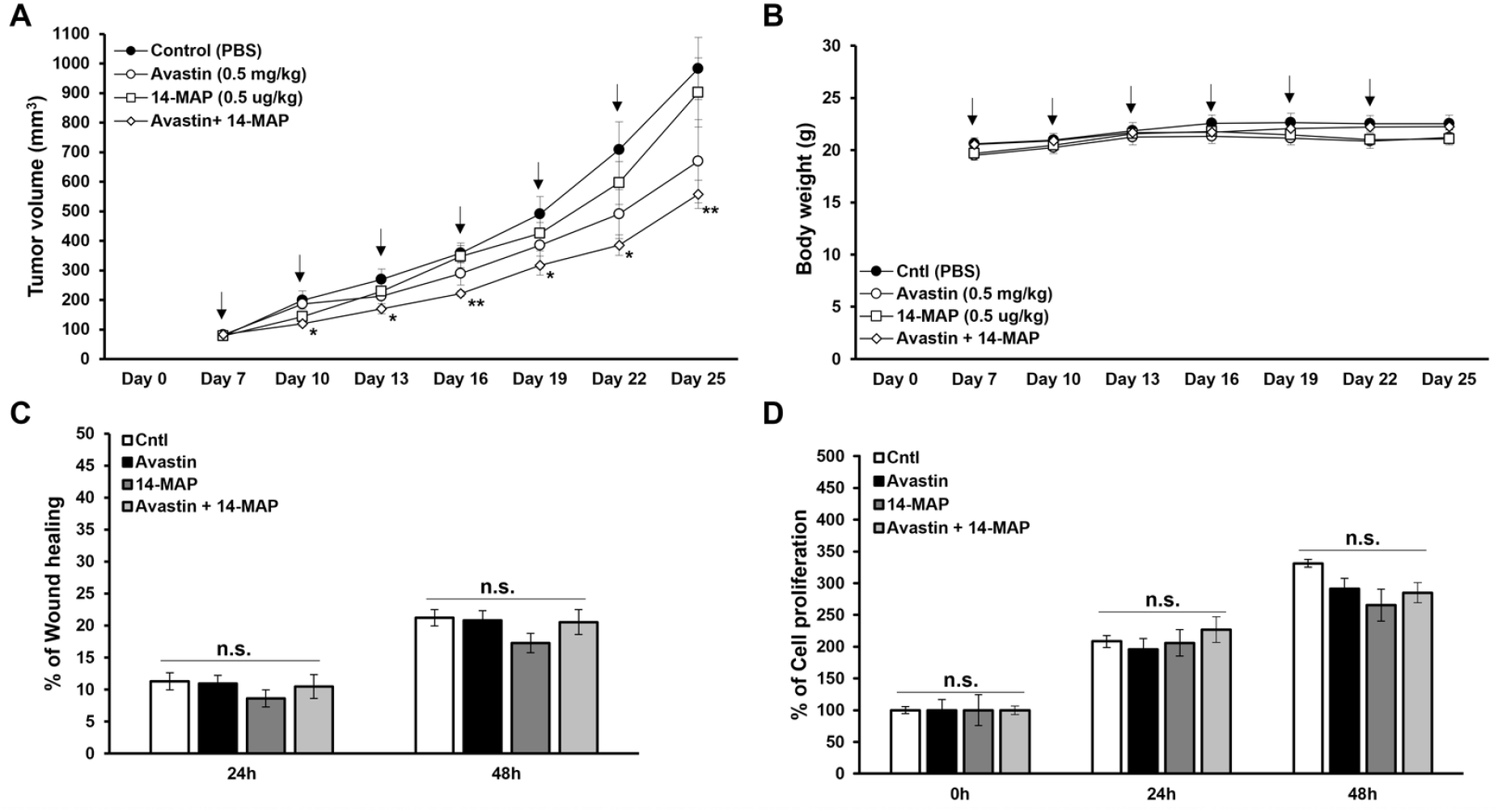
14-MAP effects on cancer. **A**, Tumor growth monitoring in HT29-xenografted Balb/c nude mice. **B**, Time-dependent body weight monitoring of the mice. **C**, HT29 cell migration analysis. **D**, HT29 cell proliferation analysis. Cntl, control (PBS). Arrows indicated I.V. injection (**A** and **B**). Avastin (M.W., 149 kDa) and 14-MAP (M.W., 1.4 kDa) were treated as 0.5 mg/kg and 0.5 ug/kg, respectively (**A** and **B**), and 5 ng/mL (approx. 34 pM) and 50 pg/mL (approx. 35 pM), respectively (**C** and **D**). *, *P* < 0.05; **, *P* < 0.01 *vs* control.

Meanwhile, the effect of 14-MAP on colon cancer cell migration and proliferation was investigated (Figure 4C and D). The treatment of 14-MAP (35 pM), Avastin (34 pM) or the combination of both showed no significant differences compared to control on cancer cell migration and cancer cell proliferation up to 48 h, but a slightly inhibited pattern was monitored after 14-MAP treatment rather than the combination treatment on both cancer cell functions (Figure 4C and D). Moreover, this 14-MAP effect on cancer cells was confirmed in the results of colony formation for a longer period (over a week) (Figure S6). In the human colon cancer cells (HT29) and human breast cancer cells (MCF-7), the treatment of 14-MAP, Avastin or the combination of both did not induce any significant difference compared to control (Figure S6A and B).

## DISCUSSION

Previously, we identified the 14-MAP sequence located at the second loop between the first and second α-helices (H1-H2) of N-terminal 14K buPRL and ascertained the antiangiogenic activity of the synthesized peptide by *in vitro* and *ex vivo* studies (8). The 14-MAP sequence was confirmed to have a specifically antiangiogenic activity *in vitro* and *ex vivo*. Compared to the negative control (Scr), randomly selected among randomly rearranged sequences, as well as the control (PBS), 14-MAP significantly inhibited EC migration (Figure 1A and B), NO production (Figure 2A) and chick vessel development (Figure S1). 14-MAP showed properties in common with its parental protein, N-terminal 14K buPRL (8).; **1)** the very low working concentration of the peptide (Figures 1-5 and Figure S2A) and **2)** its antiangiogenic function associated with VEGF- and BK-dependent eNO production (Figure 2A). We observed this kind of low concentration activity of the peptide in our previous study (25). Their highly sensitive and effective role at the picomolar concentration hints at specific targets like a high affinity binding partner, although we did not identify it in this study. Other anti-angiogenic protein and peptide candidates like a Wnt binding partner, secreted frizzle-related protein 4 showed high antiangiogenic effects at low working concentrations as low as 3 nM (25). Recently, the peptide therapeutics market has been gaining importance (34,35). Indeed, 7.2% of total drugs (15/208) approved by FDA in the last few years (2015-2019) are peptide-based drug (36). Among the peptide-based drugs, small or medium sized peptide drugs such as Macimorelin (Macrilen^®^, 3-mer), Plecanatide (Macrilen^®^, 16-mer), Afamelanotide (Scenesse^®^, 13-mer), etc. account for more than half (36). The development of peptide-based drugs has been of interest because of the advantages of size, easy delivery, specificity, and being biologic-based (34). In a similar line of thought we anticipate that 14-MAP will be a promising candidate for the peptide therapeutics market.

In the present study, we directed our attention to two well-accepted angiogenic pathways, the VEGF- and BK-mediated eNOS pathways (19,20). The study revealed that the antagonistic function of 14-MAP was on both-dependent NO-mediated effects in EC (Figure 1C-G and Figure 2A), although the function of the peptide was much more correlated to BK-dependent angiogenic functions than VEGF-dependent functions (Figure 1C, D and F). This could suggest that 14-MAP might target matured vessels such as small arteries and arteries niche within the tumor microenvironment (37). Moreover, we observed 14-MAP-mediated interference in the BK/B2R pathway more than the Des- Arg9-BK/B1R pathway in EC (Figure 3). The interference of 14-MAP-mediated BK-receptor pathway requires to identify the antagonistic route with more studies. Furthermore, we demonstrated that 14- MAP inhibited SpNO- and SC-induced NO production/EC migration of the eNOS/NO/sGC/cGMP/PKG pathway, which happens the early stage of angiogenesis (Figure 2). However, it is not clear how 14-MAP interacts with NO downstream cGMP signaling.

The potential of a BK- and eNOS-dependent antiangiogenic peptide on cancer therapy was tested (Figure 4). 14-MAP at the low dose significantly enhanced tumor reduction by Avastin treatment in human colon cancer-xenografted to nude mice (Figure 4A). The enhancing effect of 14-MAP on Avastin-reduced cancer progression *in vivo* was not monitored in *in vitro* studies (Figure 4C and D, and Figure S6). Therefore, the effect of 14-MAP on cancer treatment is considered as no cancer cell-specific. Currently, due to the development of resistance to antiangiogenic therapy, targeting VEGF-dependent angiogenesis requires to turn to targeting VEGF-independent angiogenesis (5). Although the effect of 14-MAP on the reduction of tumor progression was less than our expectation in the present study, the antiangiogenic peptide at a very low dose successively intensified the tumor reduction by the low dose of Avastin. This effect of 14-MAP on tumor reduction by Avastin is interpreted as an additive anti-tumor effect of 14-MAP, which interferes with BK signaling in endothelial cells. These results suggest that the combination therapy of Avastin and 14-MAP can overcome the limitations of VEGF-targeted antiangiogenic therapy.

Meanwhile, Bajou *et al* suggested that N-terminal 16K human PRL could work for anticancer therapy by inducing fibrinolysis as well as inhibition of angiogenesis (38). Their study confirmed plasminogen activator inhibitor-1 (PAI-1) as the direct binding partner of 16K PRL (amino acid residue 176-228 and 244-289, as binding site 1 and 2, respectively) and demonstrated that 16K PRL antagonized PAI-1 complex (PAI-1/uPA/uPAR)-induced thromboembolism and angiogenesis (38). We do not perceive that 14-MAP directly binds to PAI-1 because 14-MAP sequence maintains a distance from the two binding sites of 16K PRL. Rather, 14-MAP sequence is the part of the antiangiogenic domain of vasoinhibin (39). Recently, Robles *et al* reported the antiangiogenic domain of vasoinhibin (1-79 residues) derived from the N-terminal human PRL fragment (39). The antiangiogenic mechanism of 14-MAP probably closes to that of the domain of vasoinhibin. BK, widely expressed in vascular tissue, induces pathophysiological process including angiogenesis, inflammation, and thromboembolism (9-11). BK up-regulates PAI-1 mRNA and protein by NO-mediated process in EC (11). The thromboembolism of vessels is specified as one of the adverse effects of antiangiogenic agents, including Avastin in cancer patients (40). We found that the combination of BK-dependent antiangiogenic peptide (14-MAP) and VEGF-dependent antiangiogenic agent (Avastin) was the most efficient in reducing tumor growth *in vivo*. Our BK-dependent antiangiogenic peptide can be useful by minimizing the adverse effects of traditional VEGF-dependent antiangiogenic agents in cancer therapy. Therefore, our study supports the development of multidrug therapeutics for anticancer therapies.

Further, antiangiogenic properties of 14-MAP need in depth investigations into its relations with other hallmarks of cancer. Questions to be addressed are: is the 14-MAP efficient enough to target other forms of angiogenesis like intussusceptive angiogenesis, vascular mimicry or co-option? Since tumor-vasculature depends on the cross talk among stakeholders in tumor microenvironment like immune cells, cancer stem cells, macrophages and fibroblasts, understanding cellular and biochemical talks among all stake holders with 14-MAP sensitized tumor cells is critical for unravelling the mechanism of 14-MAP dependent inhibition of tumor angiogenesis (41-43). We envisage the further direction of research in 14-MAP based inhibition of tumor angiogenesis warrants support from omics based interactome studies of tumor microenvironment.

## AUTHOR CONTRIBUTIONS

Conceptualization, J.L., S.M., P.K.J., K.M. and S.C.; methodology, S.M., P.K.J., S.N., J.L. and S.C.; formal analysis, J.L., S.M., K.M. and S.C.; investigation, J.L., P.K.J., S.N., S.H.P., L.S., J.-S. C., H.J.L., K.M. and S.C.; resources, J.-S.C. and K.M. data curation, J.L.; writing – original draft preparation, J.L., P.K.J., H.J.L. and S.C.; writing – review & editing, J.L., P.K.J., H.J.L. and S.C.; visualization, J.L. and P.K.J.; supervision, H.J.L. and S.C.; project administration, J.L., H.J.L. and S.C.; funding acquisition, J.L., H.J.L. and S.C. All authors have read and agreed to the published version of the manuscript.

## ACKNOWLEDGMENTS

We thank Dr. J.K. Bang in KBSI, Republic of Korea for 14-MAP and Scr peptides, Dr. C.J.S. Edgel in University of North Carolina, USA for EAhy926 cells, Dr. U. Schumacher in University Medical Center Hamburger-Eppendorf, Germany for HT29 cells and Dr. V. Shah, Mayo Clinic, Rochester, USA for T24/ECV 304 eNOS-GFP cells.

## SUPPLEMENTSRY MATERIAL

The following are available online at www.mdpi.com/xxx/s1, Figure S1: No inhibitory effect of Scr on chick vessel development, Figure S2: Working concentration of 14-MAP in EC, Figure S3: 14-MAP effect on BK- and VEGF-induced EC functions, Figure S4: Diagram of BK-BKR pathways, Figure S5:In vivo analysis, Figure S6: Effect of 14-MAP on colony formation of cancer cells, Figure S7:The possibility of 14-MAP docking to BK receptors, Table S1: 14-MAP toxicity in ICR mice.

## DATA AVAILBILITY STATEMENT

Data is contained within the article or supplementary material, and the raw data supporting the conclusions of this article will be made available by the authors, without undue reservation.

