## Supplementary file for "14K Prolactin Derived 14-Mer Antiangiogenic Peptide Targets Bradykinin-/Nitric Oxide-cGMP-Dependent Angiogenesis"

### Supplementary Information

***Cell Viability Assay:*** EAhy926 cells were seeded in 24-well plates and incubated at 37 ºC and 5% $CO_2$ to achieve 60-70% confluence. The cells were treated with different concentrations of 14-MAP peptide ranging from 1 pg/mL to 10 µg/mL for 4 h (the predicted half-life (*J Biol Chem* (1989) 264:16700-16712). After treatment period, trypan blue (0.4 mg/mL) was added to the media and incubated further for 15 min. After removing media, cells were washed once with PBS and further PBS was added to the wells. The number of cells with a blue nucleus was counted manually under the 4X objective of on Olympus inverted microscope.

***Dose Determination of 14-MAP and Avastin In Vivo:*** HT29 cells (5 x $10^6$ cells/100 µL PBS and Matrigel® mixture) were subcutaneously inoculated on the flank of balb/c nude mice (5-week-old (16.3-19.5 g), female). When tumor volume was over 100 $mm^3$, two different doses of Avastin (0.5 and 5 mg/kg) and two different doses of 14-MAP (0.5 and 1 µg/kg) versus control (PBS) were given twice per week for 3 consecutive weeks by an intravenous injection to the tail vein ($n$ = 3-4/group). The tumor size was measured with a digital caliper, twice a week. The mice were euthanized when the tumor volume reached 2,000 $mm^3$ or if there was any distress before. All animal care and experimentation were followed the procedure approved by the Ewha Womans University Institutional Animal Care and Use Committee (IACUC) (No. 2016-16-062), Korea.

***Animal Toxicity Study:*** ICR mice (6-week-old (27.7-34.4 g), male) were treated with four different doses of 14-MAP (0.01, 0.1, 0.5 and 1 µg/kg) and PBS (control) every day for 8 days by an intravenous injection to the tail vein ($n$ = 5/each group). The body weight was monitored twice a week for an initial 2 weeks and then it was done once a week for the last 2 weeks. The mice were checked daily for their condition such as behavior, mortality, etc. Four weeks later, the organs such as a liver, stomach, intestines, spleen, kidney, lung, heart, etc. were ascertained in all mice. All animal care and experimentation were followed the procedure approved by the Ewha Womans University Institutional Animal Care and Use Committee (IACUC) (No. 2016-062), Korea.

***Colony Formation Assay*:** HT29 and MCF-7 cells (200/well, 24-well plate) were seeded as a single cell. After 24 h, 14-MAP (50 pg/mL), Avastin (5 ng/mL) and the combination vs control were added to the cells, and the cells were incubated to form colonies for 8-13 days. The colonies were stained with 0.5% crystal violet after methanol fixation.

### Supplementary Figures and Table

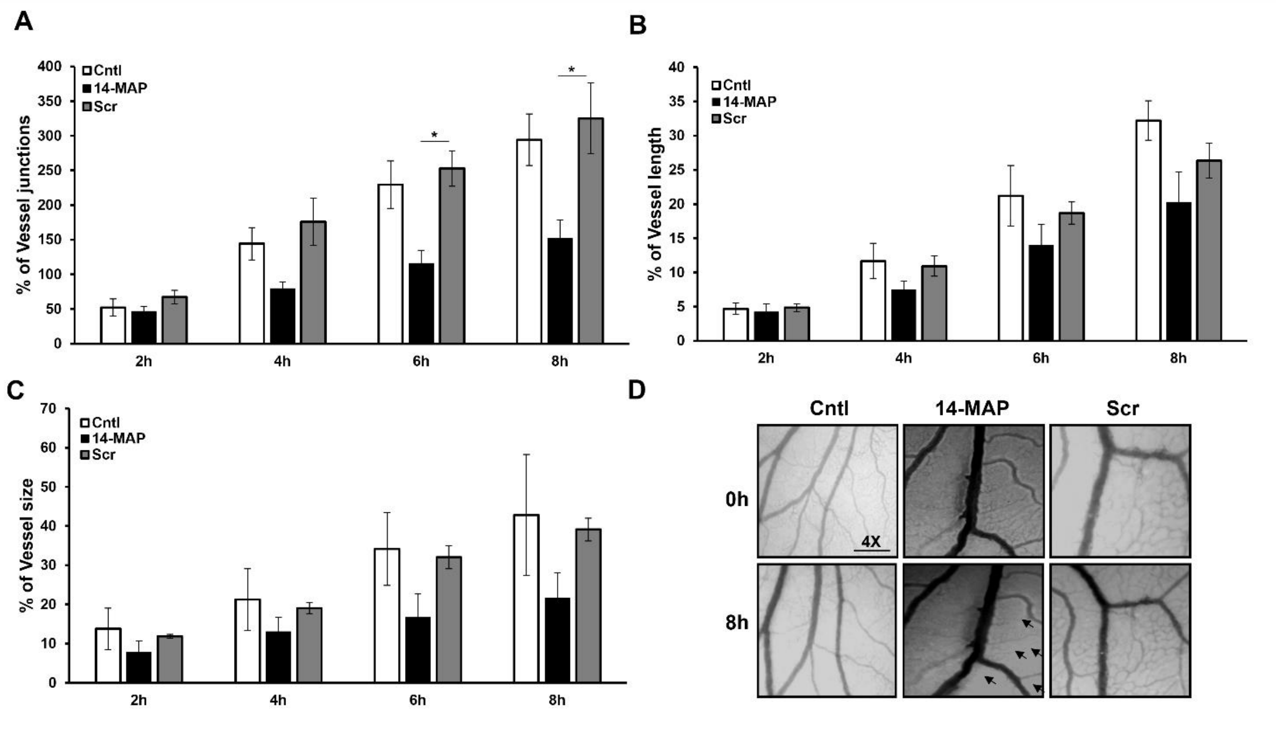


**Supplementary Figure 1. No inhibitory effect of Scr on chick vessel development.** Analysis of vessel junction (**A**), length (**B**) and size (**C**) by CAM assay. **D**, Pictures of vessel development. Cntl, control; Scr, scramble (GSQCAAGTMNLKIF). *, $P < 0.05$, arrows represent indicate the deteriorating vascular network.

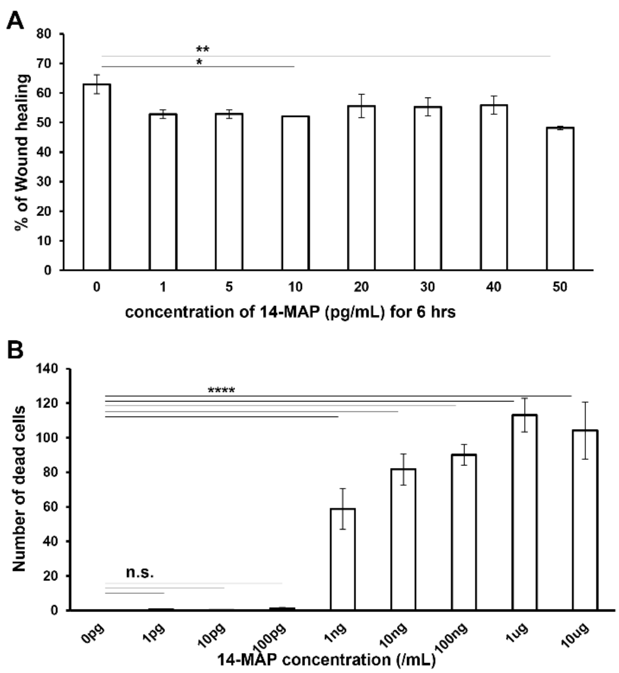


**Supplementary Figure 2. Working concentration of 14-MAP in EC. A**, Cell migration analysis and **B**, cytotoxicity analysis after treatment with 14-MAP peptide (M.W. 1.4 kDa). n.s., no significant; *, $P < 0.05$; **, $P < 0.01$; ****, $P < 0.0001$.

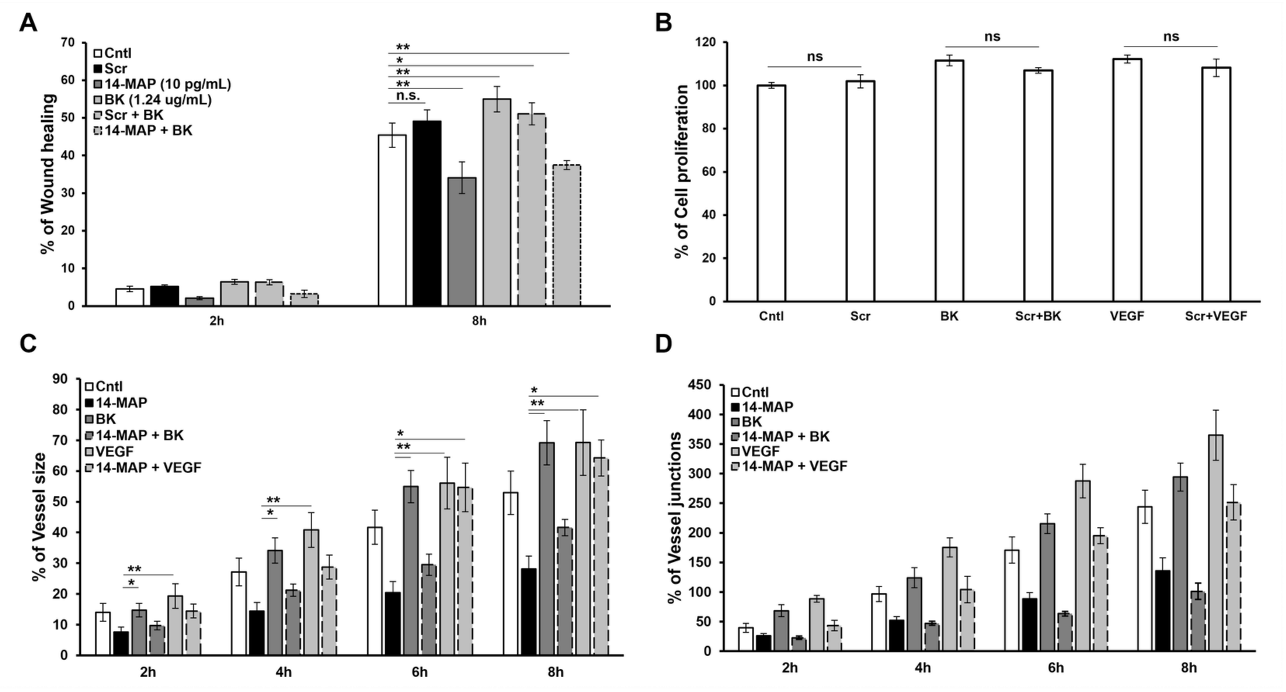


**Supplementary Figure 3. 14-MAP effect on BK- and VEGF-induced EC functions.** **A**, 14-MAP effect on BK-induced cell migration. **B**, Scr effect on BK- and VEGF-increased cell proliferation. **C** and **D**, 14-MAP effect on BK- and VEGF-improved chick vessel size and junction, respectively. Cntl, control; Scr, scramble (GSQCAAGTMNKIF). n.s., no significant; *, $P < 0.05$; **, $P < 0.01$.

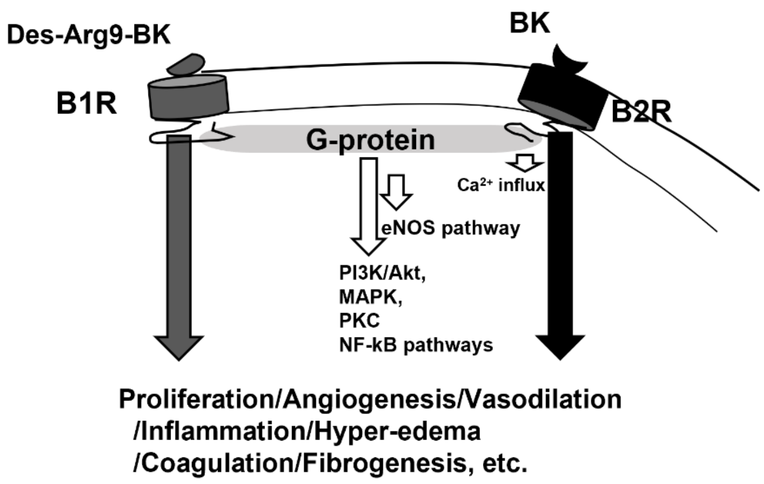


**Supplementary Figure 4. Diagram of BK-BKR pathways.**

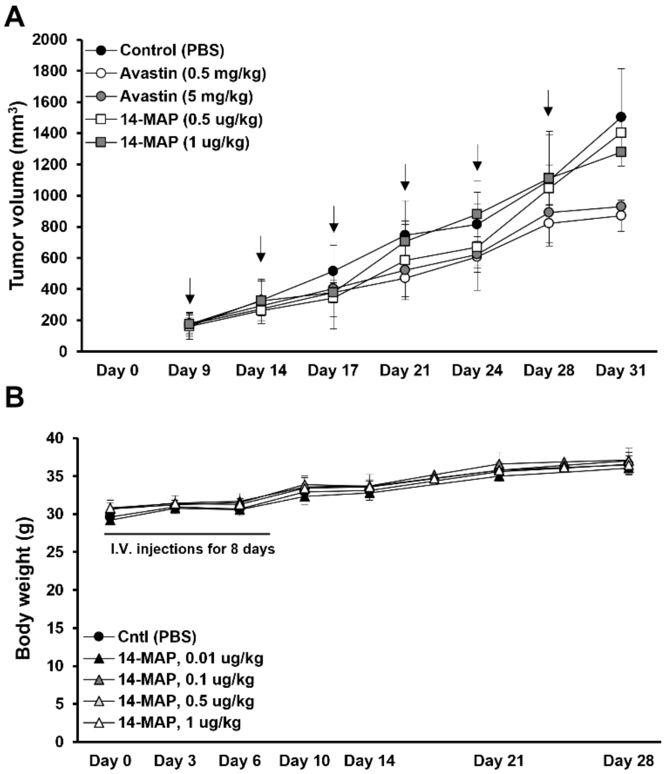


**Supplementary Figure 5.** ***In vivo* analysis. A**, Dose determination of 14-MAP (M.W., 1.4 kDa) and Avastin (M.W., 149 kDa) in Balb/c nude mice. **B**, Toxicity analysis (body weight) after 14-MAP treatment to ICR mice. Cntl, control. Arrows indicated I.V. injection.

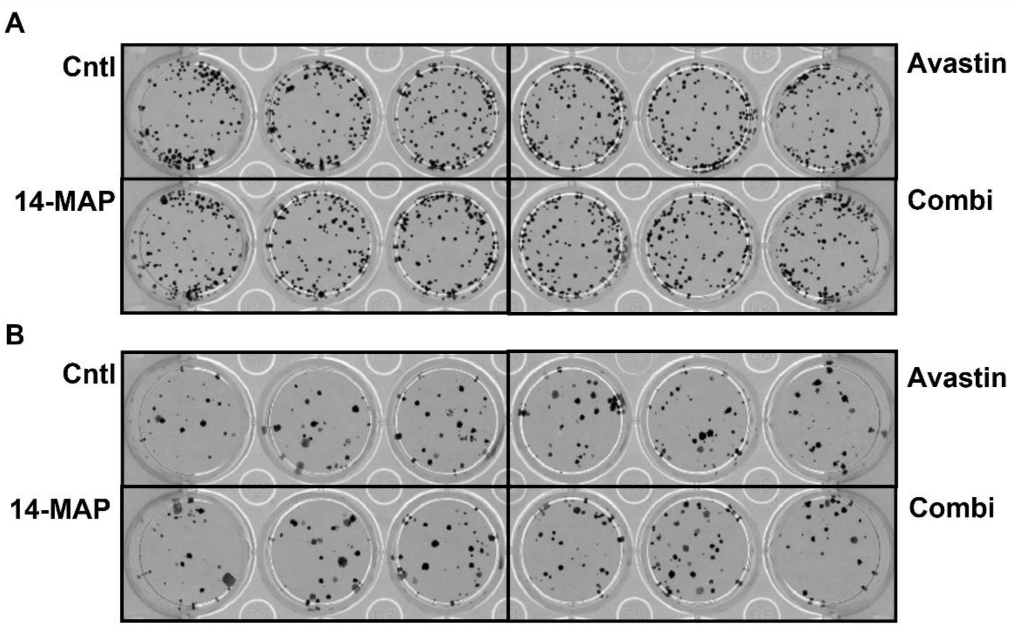


| A | Cntl | Avastin | 14-MAP | Combi | B | Cntl | Avastin | 14-MAP | Combi |
|---|---|---|---|---|---|---|---|---|---|
| | 9 | 20 | 10 | 17 | | 80 | 90 | 90 | 85 |
| | 15 | 13 | 18 | 24 | | 70 | 80 | 85 | 83 |
| | 20 | 13 | 17 | 17 | | 90 | 85 | 75 | 90 |
| Mean | 14.67 | 15.33 | 15.00 | 19.33 | Mean | 80 | 85 | 83.33 | 86.00 |
| S.D. | 5.51 | 4.04 | 4.36 | 4.04 | S.D. | 10 | 5 | 7.64 | 3.61 |

**Supplementary Figure 6. Effect of 14-MAP on colony formation of cancer cells. A**, HT29 cells and **B** MCF-7 cells. Cntl, control, PBS; Avastin, 5 ng/mL (approx. 34 pM); 14-MAP, 50 pg/mL (approx. 35 pM); combi, combination of Avastin and 14-MAP.

**Supplementary Table 1. 14-MAP toxicity in ICR mice.**

| Group | Survival Rate (%) |
|---|---|
| Control (PBS) | 100 |
| 14-MAP, 0.01 μg/kg | 100 |
| 14-MAP, 0.1 μg/kg | 100 |
| 14-MAP, 0.5 μg/kg | 100 |
| 14-MAP, 1 μg/kg | 100 |
